# MACE2K: A Text-Mining Tool to Extract Literature-based Evidence for Variant Interpretation using Machine Learning

**DOI:** 10.1101/2020.12.03.409094

**Authors:** Samir Gupta, Shruti Rao, Trisha Miglani, Yasaswini Iyer, Junxia Lin, Ahson M. Saiyed, Ifeoma Ikwuemesi, Shannon McNulty, Courtney Thaxton, Subha Madhavan

## Abstract

Interpretation of a given variant’s pathogenicity is one of the most profound challenges to realizing the promise of genomic medicine. A large amount of information about associations between variants and diseases used by curators and researchers for interpreting variant pathogenicity is buried in biomedical literature. The development of text-mining tools that can extract relevant information from the literature will speed up and assist the variant interpretation curation process. In this work, we present a text-mining tool, MACE2k that extracts evidence sentences containing associations between variants and diseases from full-length PMC Open Access articles. We use different machine learning models (classical and deep learning) to identify evidence sentences with variant-disease associations. Evaluation shows promising results with the best F1-score of 82.9% and AUC-ROC of 73.9%. Classical ML models had a better recall (96.6% for Random Forest) compared to deep learning models. The deep learning model, Convolutional Neural Network had the best precision (75.6%), which is essential for any curation task.

## 1 Introduction

The interpretation of any given variant’s pathogenicity is one of the most profound challenges to realizing the promise of genomic medicine. It is imperative to understand how gene variants impact certain diseases and associated phenotypes. Consortium projects like ClinGen [1], gnomAD [2], and GA4GH [3] are curating knowledge bases for understanding the clinical relevance of human genetic variation, based on novel methods for assessing the clinical actionability of genes and the pathogenicity of genetic variants. Literature review to identify published assertions of associations between variants and diseases is a crucial step variant interpretation curation process. There has been a rise in the volume of scientific literature describing variant-disease assertions due to sequencing techniques [4]. Manually curated databases through literature review containing variants and associated diseases such as COSMIC [5], BioMuta [6], OMIM [7], HGMD [8], dnSNP [9], ClinVar [10], CIViC [11] have been developed. It is becoming increasingly difficult for biocurators, clinical researchers, and clinicians to keep up with the rapidly growing volume and breadth of variant-related information from published literature. The value of extracting and understanding genetic variations and their relationship to disease from literature has been recognized [12] and there is a pressing need to develop text-mining tools to extract evidence of variant-disease associations from literature. Extracting such relevant information from literature will speed up and assist the manual variant interpretation curation process. To address such variant interpretation curation needs, we have developed a text-mining tool named MACE2K to extract evidence sentences indicating variant-disease associations from full-length PMC articles. We will treat this extraction task as a classification problem i.e. given a sentence with the variant and disease annotated, we will train ML models to classify the sentence as positive or negative indicating the presence or absence of the variant-disease association. We train and test different classical ML models: Logistic Regression, Support Vector Machines, Random Forest, and deep learning models: Convolutional Neural Networks (CNN) and Long short-term memory (LSTM) for evidence sentence classification. The different ML classifiers were evaluated using 5-fold cross-validation on an in-house curated dataset and achieved average precision, recall, and F1-score of up to 75.6%, 96.6%, and 82.9%, respectively.

## 2 Methods

As indicated earlier, MACE2k is a text-mining tool that extracts evidence sentences indicating variant-disease associations from full-length PMC articles. An example positive sentence indicating a variant-disease association is provided in Example 1 below. In this instance, an association between the variant “His239Arg”, the associated gene “HRAD9”, and the disease “lung adenocarcinoma” is stated.

Example 1: <u>His239Arg</u> SNP of <u>HRAD9</u> is associated with <u>lung adenocarcinoma</u>. [PMID: 16444745]

We treat this extraction task as a classification problem i.e. given a sentence with the variant and disease annotated, we will train ML models to classify the sentence as positive or negative indicating the presence or absence of the variant-disease association. We will use

PubTator [13] for entity typing i.e. to detect and normalize the gene, variant, and disease mentioned in a sentence to be classified. In a previous work, we have developed a pattern-based relation extraction system called eGARD [14] to extract such relationships between variants, disease, and drug responses from abstracts. As with any rule/pattern-based approach, eGARD suffered from low recall. In this work, we employ Machine Learning (ML) to address the issue of low recall and additionally extend the extraction to full-length PMC articles. We present an overall workflow of the study design and methodologies in Fig 1. The different steps of the approach are (1) Creation of a curated dataset, (2) Extraction of features from evidence sentences for training ML models, and (3) Training and evaluation of ML models. These steps are described in the subsequent subsections.

**Fig 1.**
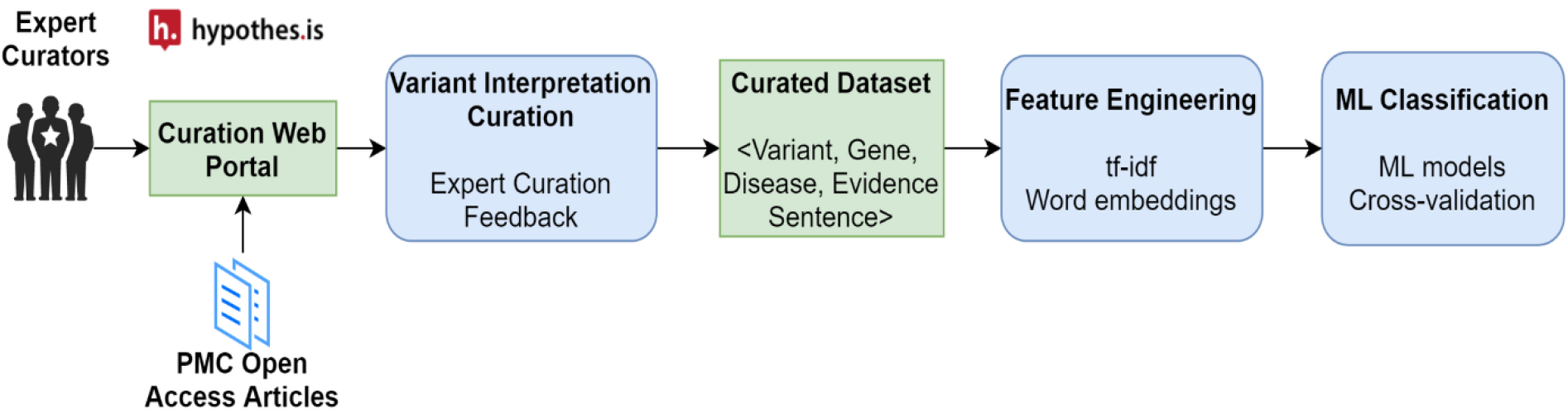
System Workflow. Abbreviations used: tf-idf (term frequency-inverse document frequency)

### 2.1 Creation of the curated dataset

For the creation of this dataset, we formulated an annotation protocol, which was provided to the curators. The object of this annotation experiment was to highlight the evidence sentences in full-length PMC articles that indicate an assertion by the author for a relationship between any pair of the three entities: (1) disease, (2) gene, and/or (3) variant. Additionally, the entities (disease, gene, variant) with normalized identifiers (HGNC [15], MONDO [16], ClinVar [10]) in the evidence sentences were also marked by the annotator. If no associated ClinVar identifier existed for the annotated variant, the curator used a ClinGen Allele Registry (CA) [17] identifier to normalize the variant. An annotation tool called Hypothes.is [18] was used to assist in the creation of the curated gold set. Based on the annotation protocol, 1000 evidence annotations (gene-variant, gene-disease, or variant-disease associations) from 87 PMC Open Access articles were annotated. The different statistics of the curated dataset are depicted in Table 1. A total of 557 evidence sentences were annotated out of which 181 sentences contained a variant-disease association. As our aim is to extract evidence sentences containing a variant-disease association relevant for variant pathogenicity interpretation, we used these 181 evidence sentences as positive instances for our ML models.

**Table 1.** Curated dataset statistics

| Category | Number of instances |
| --- | --- |
| PMC Open Access Articles | 87 |
| Gene Entities | 137 |
| Variant Entities | 186 |
| Disease Entities | 136 |
| Evidence Sentences with any entity-entity relation | 557 |
| Evidence Sentences with variant-disease relation | 181 |

Since the annotation protocol only required the curator to identify sentences, where an association between a variant and disease was asserted, all these sentences are positive instances of evidence sentences. Since negative instances are required to train ML models, we used an automated approach to identify such negative sentences. From the curated PMC open-access articles, we downloaded the associated manuscript text and relevant entities (gene, variant, disease) using PubTator’s APIs [19] in BioC-XML format. The XML output was parsed and text associated with each section in the manuscript was split into sentences using the Stanford CoreNLP toolkit [20]. For a particular curated PMC article, all sentences with a variant and disease mention (as determined by PubTator) that were not annotated by the curator assigned as negative instances. Our assumption is that a sentence where a variant and disease co-occur and not identified by the curator for a variant-disease association is not relevant for variant pathogenicity interpretation and should be assigned a negative label for ML training purposes. This approach yielded a total of 98 negative evidence sentences. Thus our curated dataset for ML training contained 181 positive and 98 negative instances.

### 2.2 Feature Engineering

In order to train ML models, the evidence sentences for positive and negative annotations need to be converted to structured features. The first set of features, which we used as input for our classical ML models is term frequency-inverse document frequency (tf-idf) weighting [21]. The second set of features, which we used as input to our deep learning models are distributed word embeddings, which have been shown to achieve better performance in NLP tasks by learning similar vectors for similar words [22,23]. For our deep learning models, we used the publicly available pre-trained word embeddings from NCBI: BioWordVec [24,25]. BioWordVec are biomedical word embeddings, trained on PubMed and clinical notes from the MIMIC-III Clinical Database [26] using fastText [27]. The word vectors have a dimensionality of 200. Words that were not present in the set of pre-trained words are set as a zero vector.

### 2.3 Classification Machine Learning Models

Different classical and deep learning ML models were employed to classify sentences indicating a variant-disease association or not. The different shallow ML models with tf-idf feature representation that we tried are Logistic Regression (LR), Support Vector Machine (SVM), and Random Forest (RF). Default parameters and loss function were used. Additionally, we used pre-trained word embeddings (BioWordVec) to build and test two deep learning architectures, namely, Convolutional Neural Network (CNN) and Bi-directional Long Short-Term Memory (bi-LSTM). The CNN architecture had a one-dimensional convolutional layer with rectified linear unit (ReLU) activation. For the convolutional layer, we experimented with three windows sizes: 3, 5, and 7, each of which has 400 filters. Every filter performs convolution on the text matrix and generates variable-length feature maps. We got the best results with a single-window of size 5. Global max pooling was then performed over each map, which essentially extracts fixed-length global features for the text. A dense layer of size 1 with a sigmoid activation function was applied over the global features to obtain the CNN classifier. The biLSTM architecture consisted of a 64-cell bidirectional LSTM layer followed by two pooling layers (maximum and average). The maximum and average pooling layers were concatenated fed to a fully connected layer with 64 units (with ReLU activation). A dense layer of size 1 with a sigmoid activation function was applied over the fully connected layer to obtain the biLSTM classifier. For the deep learning models, we used binary cross-entropy as the objective loss function and the Adam algorithm [28] to optimize the loss function. To train the parameters for CNN and bi-LSTM, we used mini-batch training with a batch size of 32. In between each layer of our deep learning architectures, we added a dropout layer with a dropping probability of 0.5 to avoid overfitting during training. We set the number of epochs to 100 during training.

## 3 Results and Discussion

We compared the performance of the different ML models to classify evidence sentences using 5-fold cross-validation. Average precision, recall, F1-score, and area under the receiver operating characteristic Curve (AUC-ROC) were computed across the folds and reported in Table 2 below. SVM had the best performance in terms of F1-score 82.9% and an AUC-ROC of 73.9%. Classical ML models had a better recall (best 96.6% for Random Forest) compared to deep learning models. The deep learning model CNN has the best precision of 75.6%.

**Table 2.** Evaluation results

| Model | Precision | Recall | F1-score | AUC-ROC |
| --- | --- | --- | --- | --- |
| **LR** | 68.1 | 95.4 | 79.5 | 63.6 |
| **SVM** | 74.1 | 94.3 | **82.9** | **73.9** |
| **RF** | 71.1 | **96.6** | 81.8 | 60.1 |
| **CNN** | **75.6** | 84.7 | 79.9 | 73.7 |
| **biLSTM** | 70.2 | 78.4 | 73.7 | 67.9 |

An initial analysis of the errors made by the deep learning models was conducted. We observed that some of the negative instances in our dataset that were generated automatically were incorrect. As indicated earlier, we assigned a negative label to any sentence with variant and disease mention that was not annotated by a curator. Our approach will incorrectly label a positive sentence (with a variant-disease association) that was missed during the annotation process as a negative instance. Subsequent verification of the automatically labeled negative instances by a curator will resolve this issue.

A potential threat to the validity of our results is the small set of evidence annotation sentences (181 positive and 98 negative) with a variant and disease mention. We plan to add more curations to our dataset to validate and additionally improve the results. Although deep learning models can achieve good performance without complex human-engineered features, it requires large amounts of data to effectively train the numerous parameters in the model. As a future step, we aim to investigate various state-of-the-art ML techniques to learn from a small gold set and a large “noisy” automatically labeled data set with approaches such as transfer learning [29,30], distant supervision [31–33], and adversarial networks [34,35]. We will automatically generate large amounts of distantly labeled data using existing knowledge bases with known variant-disease entity pairs such as CIViC [11], ClinGen [1], and ClinVar [10]. Noise-reduction heuristics will be used to remove noise in the distantly-labeled data. We will first train our models on the large distantly labeled set and then re-train the model on the “small” amount of human-labeled data to increase accuracy. The intuition behind this two-step training is that an ML model trained on a large distantly labeled data is a better starting point in terms of learning the model parameters than only training on the small human-labeled data. This idea is similar to transfer learning, which leverages labeled data in a different (but similar) domain to help training models in the domain of interest that does not have sufficient training data.

## 4 Conclusion

Information regarding the evidence of variant and disease associations is largely buried in biomedical literature. This information is critical for determining variant pathogenicity and its clinical utility. It is becoming difficult for researchers and curators to keep up with the increasing amount of published literature about variant-disease associations. We have developed a textmining tool, MACE2K to extract evidence sentences with variant-disease associations from fulllength PMC Open Access articles using machine learning (ML). We compare the performance of different ML models (classical and deep learning) with different feature representations (frequency-based, distributed word embeddings) for classifying sentences containing variant-disease association or not. Our results indicate SupportVector Machines performs the best overall (F1-score: 82.9 %) for evidence sentence classification, while Convolutional Neural Network (CNN) has the best precision (75.6%), which is essential for any downstream curation task. We believe that MACE2k will assist and speed up the variant interpretation curation process by extracting relevant information from the literature.

## Notes

### Competing Interest Statement

The authors have declared no competing interest.

## References

1. Rehm HL, Berg JS, Brooks LD, Bustamante CD, Evans JP, Landrum MJ, et al. ClinGen--the Clinical Genome Resource. N Engl J Med. 2015;372: 2235–2242.

2. Karczewski KJ, Francioli LC, Tiao G, Cummings BB, Alföldi J, Wang Q, et al. The mutational constraint spectrum quantified from variation in 141,456 humans. Nature. 2020;581: 434–443.

3. Rahimzadeh V, Dyke SOM, Knoppers BM. An International Framework for Data Sharing: Moving Forward with the Global Alliance for Genomics and Health. Biopreserv Biobank. 2016;14: 256–259.

4. Zhang J, Chiodini R, Badr A, Zhang G. The impact of next-generation sequencing on genomics. Journal of Genetics and Genomics. 2011. pp. 95–109. doi:10.1016/j.jgg.2011.02.003

5. Forbes SA, Bindal N, Bamford S, Cole C, Kok CY, Beare D, et al. COSMIC: mining complete cancer genomes in the Catalogue of Somatic Mutations in Cancer. Nucleic Acids Res. 2011;39: D945–50.

6. Wu T-J, Shamsaddini A, Pan Y, Smith K, Crichton DJ, Simonyan V, et al. A framework for organizing cancer-related variations from existing databases, publications and NGS data using a High-performance Integrated Virtual Environment (HIVE). Database. 2014;2014. doi:10.1093/database/bau022

7. Amberger J, Bocchini CA, Scott AF, Hamosh A. McKusick’s Online Mendelian Inheritance in Man (OMIM®). Nucleic Acids Res. 2008;37: D793–D796.

8. Stenson PD, Mort M, Ball EV, Howells K, Phillips AD, Thomas NS, et al. The Human Gene Mutation Database: 2008 update. Genome Med. 2009;1: 13.

9. Sherry ST, Ward MH, Kholodov M, Baker J, Phan L, Smigielski EM, et al. dbSNP: the NCBI database of genetic variation. Nucleic Acids Res. 2001;29: 308–311.

10. Landrum MJ, Lee JM, Benson M, Brown G, Chao C, Chitipiralla S, et al. ClinVar: public archive of interpretations of clinically relevant variants. Nucleic Acids Res. 2016;44: D862–8.

11. Griffith M, Spies NC, Krysiak K, McMichael JF, Coffman AC, Danos AM, et al. CIViC is a community knowledgebase for expert crowdsourcing the clinical interpretation of variants in cancer. Nat Genet. 2017;49: 170–174.

12. Zhu F, Patumcharoenpol P, Zhang C, Yang Y, Chan J, Meechai A, et al. Biomedical text mining and its applications in cancer research. J Biomed Inform. 2013;46: 200–211.

13. Wei C-H, Kao H-Y, Lu Z. PubTator: a web-based text mining tool for assisting biocuration. Nucleic Acids Res. 2013;41: W518–22.

14. Mahmood ASMA, Rao S, McGarvey P, Wu C, Madhavan S, Vijay-Shanker K. eGARD: Extracting associations between genomic anomalies and drug responses from text. PLoS One. 2017;12: e0189663.

15. Gray KA, Yates B, Seal RL, Wright MW, Bruford EA. Genenames.org: the HGNC resources in 2015. Nucleic Acids Res. 2015;43: D1079–85.

16. Ontology Xref Service. Mondo Disease Ontology < Ontology Lookup Service < EMBL-EBI. [cited 3 Dec 2020]. Available: https://www.ebi.ac.uk/ols/ontologies/mondo

17. Pawliczek P, Patel RY, Ashmore LR, Jackson AR, Bizon C, Nelson T, et al. ClinGen Allele Registry links information about genetic variants. Hum Mutat. 2018;39: 1690–1701.

18. dwhly. Home : Hypothesis. [cited 27 Oct 2020]. Available: https://web.hypothes.is/

19. PubTator Central API - NCBI - NLM - NIH. [cited 3 Dec 2020]. Available: https://www.ncbi.nlm.nih.gov/research/pubtator/api.html

20. Manning CD, Surdeanu M, Bauer J, Finkel JR, Bethard S, McClosky D. The Stanford CoreNLP natural language processing toolkit. Proceedings of 52nd annual meeting of the association for computational linguistics: system demonstrations. 2014. pp. 55–60.

21. Salton G, Buckley C. Term-weighting approaches in automatic text retrieval. Information Processing & Management. 1988. pp. 513–523. doi:10.1016/0306-4573(88)90021-0

22. Mikolov T, Chen K, Corrado G, Dean J. Efficient Estimation of Word Representations in Vector Space. arXiv [cs.CL]. 2013. Available: http://arxiv.org/abs/1301.3781

23. Mikolov T, Sutskever I, Chen K, Corrado GS, Dean J. Distributed Representations of Words and Phrases and their Compositionality. In: Burges CJC, Bottou L, Welling M, Ghahramani Z, Weinberger KQ, editors. Advances in Neural Information Processing Systems 26. Curran Associates, Inc.; 2013. pp. 3111–3119.

24. Zhang Y, Chen Q, Yang Z, Lin H, Lu Z. BioWordVec, improving biomedical word embeddings with subword information and MeSH. Sci Data. 2019;6: 52.

25. Chen Q, Peng Y, Lu Z. BioSentVec: creating sentence embeddings for biomedical texts. arXiv [cs.CL]. 2018. Available: http://arxiv.org/abs/1810.09302

26. Johnson AEW, Pollard TJ, Shen L, Lehman L-WH, Feng M, Ghassemi M, et al. MIMIC-III, a freely accessible critical care database. Sci Data. 2016;3: 160035.

27. fastText. [cited 12 Aug 2019]. Available: https://fasttext.cc/

28. Kingma DP, Ba J. Adam: A Method for Stochastic Optimization. arXiv [cs.LG]. 2014. Available: http://arxiv.org/abs/1412.6980

29. Corbett P, Boyle J. Improving the learning of chemical-protein interactions from literature using transfer learning and specialized word embeddings. Database. 2018. doi:10.1093/database/bay066

30. Peng Y, Yan S, Lu Z. Transfer Learning in Biomedical Natural Language Processing: An Evaluation of BERT and ELMo on Ten Benchmarking Datasets. arXiv [cs.CL]. 2019. Available: http://arxiv.org/abs/1906.05474

31. Zhang C. DeepDive: a data management system for automatic knowledge base construction. University of Wisconsin-Madison, Madison, Wisconsin. 2015. Available: http://citeseerx.ist.psu.edu/viewdoc/download?doi=10.1.1.699.5115&rep=rep1&type=pdf

32. Li G, Wu C, Vijay-Shanker K. Noise reduction methods for distantly supervised biomedical relation extraction. BioNLP 2017. 2017. pp. 184–193.

33. Su P, Li G, Wu C, Vijay-Shanker K. Using distant supervision to augment manually annotated data for relation extraction. PLoS One. 2019;14: e0216913.

34. Rios A, Kavuluru R, Lu Z. Generalizing biomedical relation classification with neural adversarial domain adaptation. Bioinformatics. 2018;34: 2973–2981.

35. Su P, Vijay-Shanker K. Adversarial Learning for Supervised and Semi-supervised Relation Extraction in Biomedical Literature. arXiv [cs.CL]. 2020. Available: http://arxiv.org/abs/2005.04277

